# Fast FEM-based Electric Field Calculations for Transcranial Magnetic Stimulation

**DOI:** 10.1101/2024.12.13.628139

**Authors:** Fang Cao, Kristoffer Hougaard Madsen, Torge Worbs, Oula Puonti, Hartwig Roman Siebner, Arno Schmitgen, Patrik Kunz, Axel Thielscher

**Affiliations:** Danish Research Centre for Magnetic Resonance, Department of Radiology and Nuclear Medicine, Copenhagen University Hospital - Amager and Hvidovre, Copenhagen, Denmark; Magnetic Resonance Section, Department of Health Technology, Technical University of Denmark, Kgs. Lyngby, Denmark; Section for Cognitive Systems, Department of Applied Mathematics and Computer Science, Technical University of Denmark, Kgs. Lyngby, Denmark; Localite GmbH, Bonn, Germany; Department of Neurology, Copenhagen University Hospital - Bispebjerg and Frederiksberg, Copenhagen, Denmark; Institute for Clinical Medicine, Faculty of Health and Medical Sciences, University of Copenhagen, Copenhagen, Denmark

**Keywords:** Transcranial magnetic stimulation, Electric field calculations, Finite-Element Method, Neuronavigation

## Abstract

**Objective:** To provide a Finite-Element Method (FEM) for rapid, repeated evaluations of the electric field induced by transcranial magnetic stimulation (TMS) in the brain for changing coil positions.

**Approach:** Previously, we introduced a first-order tetrahedral FEM enhanced by super- convergent patch recovery (SPR), striking a good balance between computational efficiency and accuracy (Saturnino *et al* 2019 *J. Neural Eng*. **16** 066032). In this study, we refined the method to accommodate repeated simulations with varying TMS coil position. Starting from a fast direct solver, streamlining the pre- and SPR-based post-calculation steps using weight matrices computed during initialization strongly improved the computational efficiency. Additional speedups were achieved through efficient multi-core and GPU acceleration, alongside the optimization of the volume conductor model of the head for TMS.

**Main Results:** For an anatomically detailed head model with ∼4.4 million tetrahedra, the optimized implementation achieves update rates above 1 Hz for electric field calculations in bilateral gray matter, resulting in a 60-fold speedup over the previous method with identical accuracy. An optimized model without neck and with adaptive spatial resolution scaled in dependence to the distance to brain grey matter, resulting in ∼1.9 million tetrahedra, increased update rates up to 10 Hz, with ∼3% numerical error and ∼4% deviation from the standard model. Region-of-interest (ROI) optimized models focused on the left motor, premotor and dorsolateral prefrontal cortices reached update rates over 20 Hz, maintaining a difference of <4% from standard results. Our approach allows efficient switching between coil types and ROI during runtime which greatly enhances the flexibility.

**Significance:** The optimized FEM enhances speed, accuracy and flexibility and benefits various applications. This includes the planning and optimization of coil positions, pre-calculation and training procedures for real-time electric field simulations based on surrogate models as well as targeting and dose control during neuronavigated TMS.

## Introduction

Rapid repeated simulations of the electric field induced by transcranial magnetic stimulation (TMS) in the brain for changing coil positions can benefit several applications, ranging from planning procedures for optimized targeting prior to the intervention [1] to online electric field visualizations during neuronavigated TMS [2]. Standard approaches based on Finite- or Boundary-Element Methods (FEM, BEM) and personalized volume conductor models derived from structural magnetic resonance images (MRI) achieve good numerical accuracy [3–5], but have been slow for the above-mentioned applications, making them time consuming or even infeasible. On the other hand, alternatives such as spherical head models, which are available in some commercial neuronavigation systems and for which computationally efficient analytical solutions exist [6,7], give only coarse estimates of the field distribution induced in the individual head [8].

Recently, several approaches have been suggested which are based on the training of surrogate models to enable fast and accurate TMS field calculations [2,9–15]. However, the practical usability of these approaches is often compromised to achieve a sufficient speed up. For example, depending on the specific approach, evaluation of the electric field might be feasible only in a predefined region-of-interest (ROI), for a specific coil type or for a predefined range of coil positions and tilts (such as assuming perfectly tangential coil placements). A recent elegant approach based on reciprocity and Huygens’ principles resolves most of these limitations, but still requires time-consuming pre-calculations for each new patient dataset and relies on GPU-based parallel computations to achieve acceleration [15].

This points to the benefits of accelerating “general-purpose” FEM- or BEM-based approaches to make them directly usable for some of these applications or to speed up the training and preparation times of approaches relying on surrogate models. In a previous study, we validated a first-order tetrahedral FEM and showed that it in combination with super-convergent patch recovery (SPR) for postprocessing of the calculated electric fields achieves a good balance between computational efficiency and accuracy [3]. Here, we strongly optimize this approach for repeated evaluations of TMS-induced electric fields for different coil positions. After detailing the optimizations, we validate the numerical accuracy of the improved method and demonstrate that it performs on par with our previous method while being substantially faster. We also demonstrate that optimizing the head model further accelerates the FEM calculations with only a moderate reduction in accuracy, enabling visualizations of the TMS-induced electric fields during neuronavigation.

## Material and Methods

### FEM implementation

A previously validated implementation of a 1^st^-order tetrahedral FEM combined with the superconvergent patch recovery (SPR) procedure served as starting point [3,16,17]. SPR acts as a postprocessing step to increase the numerical accuracy of the electric field values compared to the original FEM results, optimizing the interpolation of the electric field to any desired position in the head mesh. Here, we improved the computational efficiency of the approach for repeated simulations with the same head mesh and one or more regions-of-interest where the electric field is evaluated.

In short, for each new coil position, the FEM solves a large sparse linear system

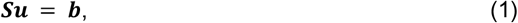

whereby ***S*** denotes the stiffness matrix depending on the head, vector ***b*** is the right-hand side (RHS) depending on the *∂**A**/∂**t*** field of the coil (the time derivative of the magnetic vector potential) and vector ***u*** denotes the unknown electric potentials at the mesh nodes. After solving for ***S***, the induced electric field ***E*** is calculated at each tetrahedra barycenter by estimating the negative spatial gradient ∇***S*** of the potential and subtracting the *∂**A**/∂**t*** field:

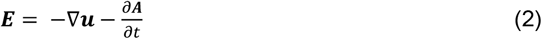

Lastly, SPR is applied to improve the numerical accuracy when interpolating ***E*** to the positions of the ROIs [18]. Details are covered in [3] and Figure 1A outlines the calculation steps. In the optimized approach described here, all parts that do not depend on the coil position are pre-calculated at startup and stored in memory (marked in green in Fig. 1A), substantially enhancing the computational efficiency for repeated simulations.:

**Figure 1.**
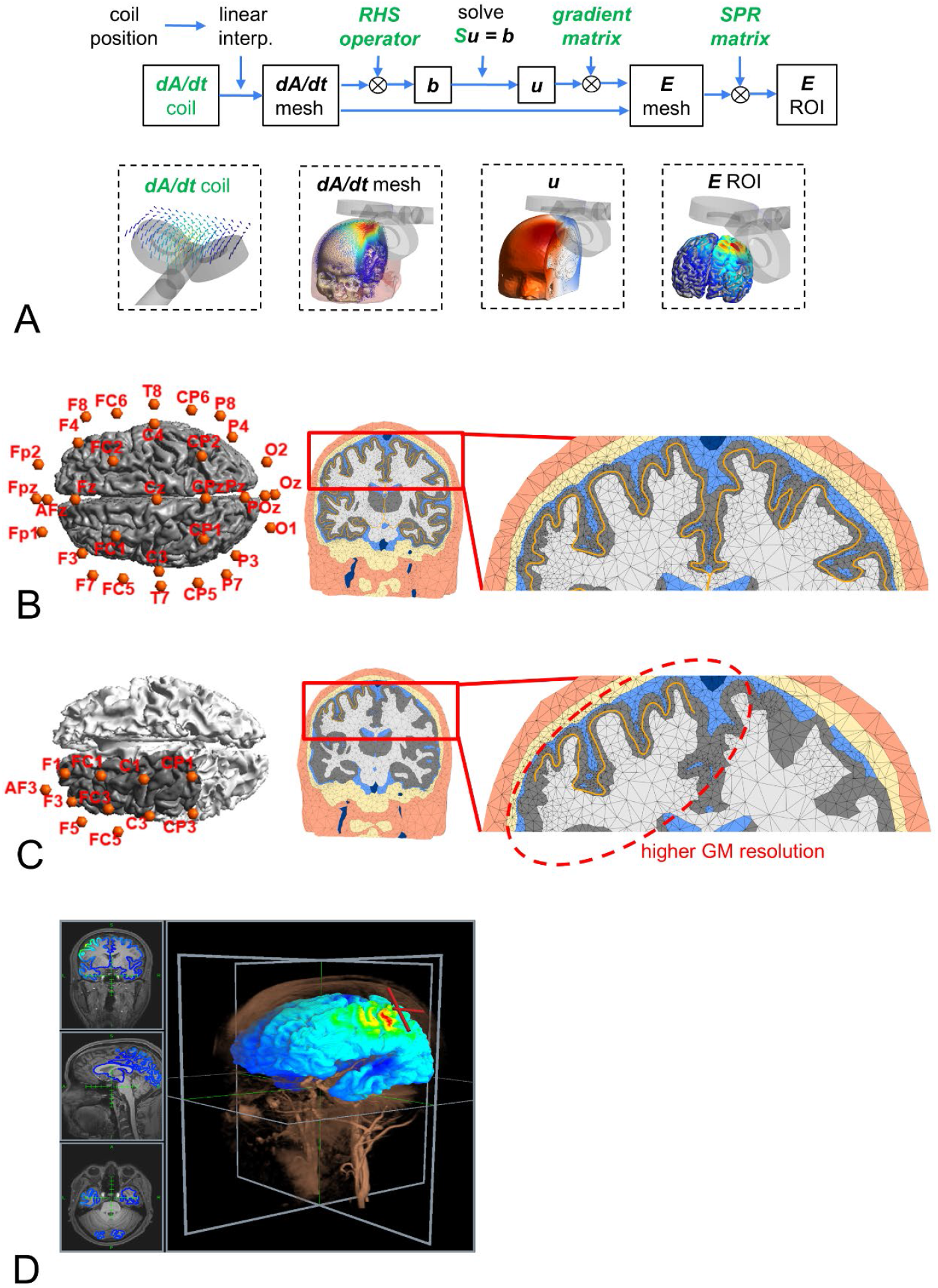
**(A)** Schematic of the calculation steps for determining the electric field for a new coil position. Items marked in green are pre-calculated and stored in memory: First, as the ***dA/dt*** field is not influenced by the head, it is calculated on a regular 3D grid for a standardized coil position once and stored as NifTI file. During runtime, the ***dA/dt*** field of a new coil position is then determined using a computationally efficient linear interpolation. Second, the stiffness matrix ***S*** is pre-calculated and the preparation step of the solver (that only requires ***S***) is run once at startup. Third, for a new coil position, the steps to determine ***b*** and ***E*** and the SPR calculations can be written as sparse matrix multiplications, whereby the matrix weights do not depend on the coil position and are thus pre-calculated at startup (⊗ denotes multiplication with a sparse matrix in compressed sparse row format). **(B)** Left: Bilateral central gray matter ROI (∼260.000 vertices, 1885cm^2^) used for calculation and visualization of the ***E***-fields, and the 32 coil positions used for testing (each with 4 orientations). The positions are chosen as subset of the EEG10-10 system. Middle and right: Optimized head mesh that has high resolution around the gray matter and high anatomical accuracy of the boundaries between gray matter, white matter, CSF and skull. **(C)** Left: Left M1+DLPFC ROI (shown in gray, ∼28.000 vertices, 210cm^2^) and the 11 coil positions used for testing (each with 4 orientations). Middle and right: Optimized head mesh with high resolution and high anatomical accuracy within 25 mm distance to the visualization ROI (indicated as orange line), and a coarser resolution in the rest of the volume. **(D)** Example of an ***E***-field visualization in the Localite TMS Navigator software.

- The *∂**A**/∂**t*** field of the coil is not influenced by the head, so that it can be pre-calculated once on a regular grid. Determining *∂**A**/∂**t*** on the mesh for a new coil position is then feasible by a combination of computationally efficient rigid transformation and interpolation. This part of the speed-up was already used in our prior implementation.
- The RHS ***b*** is calculated via a weighted summation of the *∂**A**/∂**t*** values, whereby the weights depend only on the head and are pre-calculated at startup.
- The stiffness matrix ***S*** depends only on the volume conductor model and is built and pre- conditioned at startup.
- The gradient operation used to determine ***E*** at the tetrahedra barycenters from *u* at the nodes and the SPR interpolation of ***E*** to the final positions of the ROIs are formulated as sparse matrix multiplications, whereby the matrix weights are pre-calculated at startup.

The execution speed of these calculation steps was further optimized by leveraging the multi-core architectures of modern CPUs and (optionally) GPUs. The direct solver PARDISO (version 2025.0.0 from Intel MKL) was used for the efficient parallelized solving of the sparse linear system. The pre- and post-processing steps to determine *∂**A**/∂**t***, ***b*** and ***E*** for each new coil position were implemented in custom python code using Numba (version 0.60.0) [19] that is automatically compiled at runtime to parallelized machine code using LLVM (https://llvm.org). In addition, the pre- and post-processing steps were implemented using CuPy (version 13.3.0) [20] to optionally benefit from GPU acceleration.

### Volume conductor modeling optimized for TMS

The head models created from MRI images via the default charm pipeline (complete head anatomy reconstruction method) of SimNIBS 4.1 are also aimed at electric field calculations for transcranial electric stimulation (TES). Among others, this entails modelling the skin surface via a dense triangle surface and differentiating between spongy and compact bone to enable detailed calculations of the current flow from the TES electrodes through the scalp and skull into the cranial cavity. In addition, the upper neck is represented to enable accurate simulations of low electrode positions. As these anatomical details outside the cranial cavity have little influence on the TMS-induced electric fields, we here optimized the head model to selectively reduce the number of tetrahedra. All soft tissues outside the cranial cavity were combined into a “scalp” tissue class, and spongy and compact bone into a “skull” compartment. The mesh was restricted to extend only slightly below the lowest point of the cranial cavity, whereby a suited cut-off position was determined as described further below. Finally, the mesh density (i.e., the inverse of the tetrahedra sizes) was optimized to be high in brain gray matter and the directly surrounding intracranial volumes, while being coarse in the scalp, the outer part of the skull and in deep regions of white matter and the ventricles. Compared to the head mesh of the SimNIBS example dataset “ernie” created by the default charm pipeline, the optimized mesh was reduced to a size of ∼1.9 compared to ∼4.4 million tetrahedra (Fig. 1B). The bilateral central gray matter ROI (∼260.000 vertices, 1885cm^2^) situated halfway between the pial surface and the gray-white matter boundary and on which the electric field was evaluated is indicated as an orange line. Overall, creation of the optimized head model with the adapted charm pipeline took ∼2 hrs on a laptop with 4 CPU cores (Windows 10, Intel i7-6700HQ, 32GB) which is comparable to the normal runtime of charm. Meshing used the 3D tetrahedral mesh generation of CGAL [21] and the MMG3D remeshing tools (version 5.7.0) embedded in the charm pipeline of SimNIBS 4.1.

When requiring the electric field only in a spatially restricted ROI rather than the complete gray matter, the mesh could be optimized further to have a high mesh density inside the cranial cavity around the visualization ROI (here: within 25 mm distance to the ROI), while generally reducing the density outside that volume. For a ROI covering the left motor, premotor and dorsolateral cortices (∼28.000 vertices, 210cm^2^, Fig. 1C), the resultant mesh had ∼0.9 million tetrahedra and took ∼15 min to create on the laptop.

The TMS-optimized meshes (Fig. 1B&C) included six tissue types, corresponding to white matter (WM, white, conductivity: 0.126 S/m), gray matter (GM, gray, 0.276 S/m), CSF (blue, 1.654 S/m), skull (beige, 0.01 S/m), scalp (skin tone, 0.465 S/m), and large blood vessels (dark blue, 0.6 S/m). Air cavities (e.g. the frontal sinuses) were spared in the mesh, and thereby act as non-conducting vacuums. The orange lines in Fig. 1B&C indicate the ROI positions. In the standard “ernie” dataset, the skull is instead modelled by spongy bone (0.025 S/m) and compact bone (0.008 S/m), and the vitreous bodies of the eyes (0.50 S/m) are additionally represented.

### Evaluation of Accuracy

For the TMS-optimized head meshes, we assessed the effects of changing the level of represented anatomical detail and mesh density on the accuracy of the simulated electric fields using the error metric

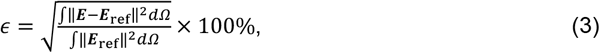

where ***E*** and ***E***_ref_ denote the electric fields calculated in the ROI for the optimized and a reference mesh respectively. Simulations were performed for 32 coil positions and the bilateral central gray matter ROI. For each position, the model of a Magstim 70mm figure-8 coil was placed tangentially on the scalp and 4 orientations were tested.

We first assessed the impact of removing the upper neck from the head model. The SimNIBS example dataset “ernie” was remeshed to have a homogenously high resolution in the lower part of the head including the upper neck, otherwise using the tissue-dependent resolution of the TMS- optimized head mesh (Fig. 2A, right panel). By deleting the corresponding tetrahedra, the mesh was then cut using axial planes at various levels from about 0.5 cm above the lowest part of the cranial cavity to 5 cm below that point. For each cutting level, the electric field in the bilateral gray matter ROI was compared to the results for the non-cut head model. These results were used to determine the caudal extent of the TMS-optimized head model for further testing.

**Figure 2.**
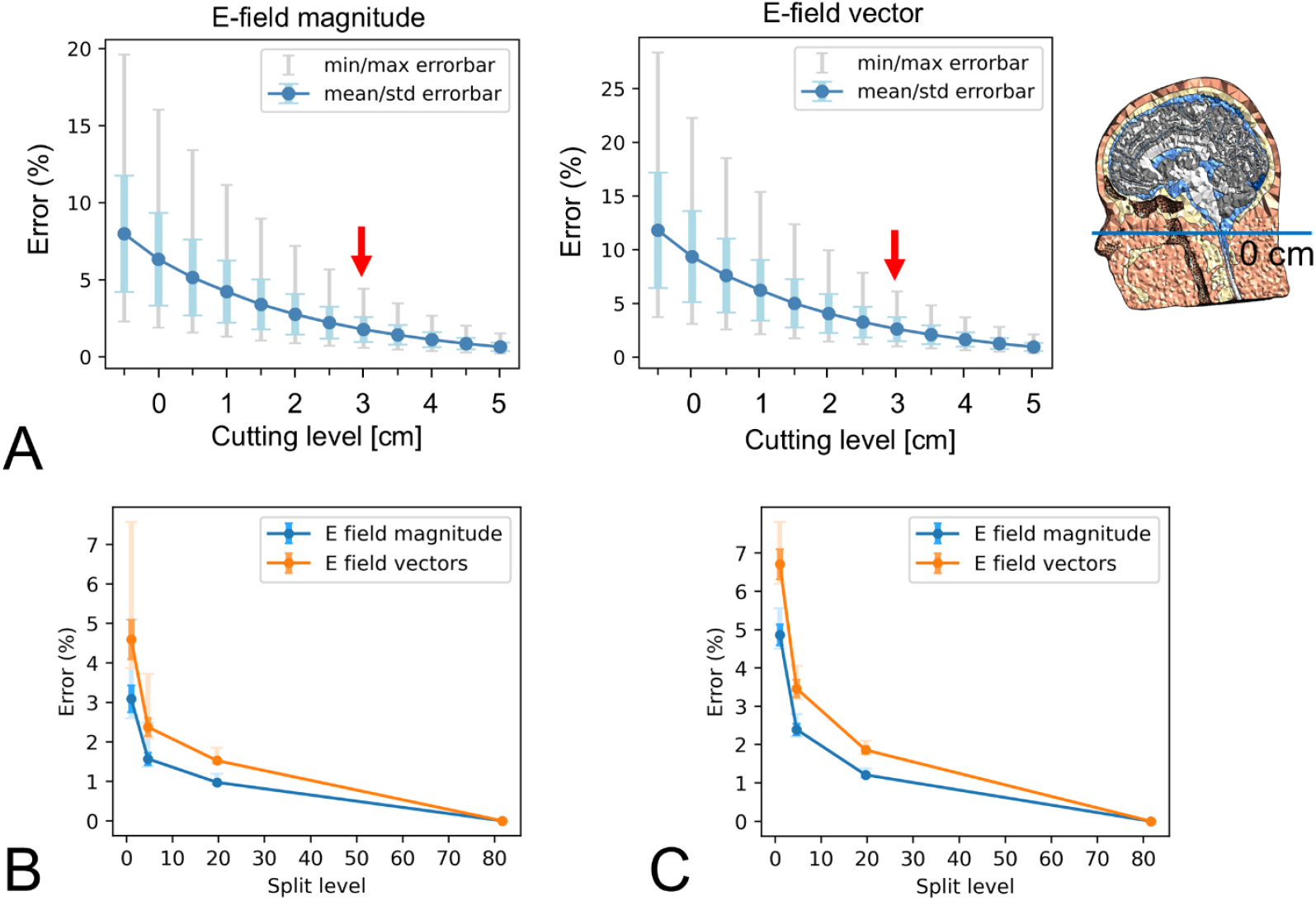
**(A)** Error levels when removing parts of the neck, relative to simulations with the non-cut mesh. The red arrows indicate the cutting level used for the TMS-optimized head meshes for the remaining results. On the right, the blue horizontal line indicates the level of the lowest part of the cranial cavity. Cuts were performed from 0.5 cm above to 5 cm below this level. **(B)** Results of the convergence analysis for the head model with ∼1.9 M tetrahedra, the bilateral central gray matter ROI and the coil positions shown in Fig. 1B. The “split level” on the x-axis indicates the factor by which the number of tetrahedra was increased, and ranges from 1 (original mesh with ∼1.9 M tetrahedra) to 83.7 (reference mesh with ∼159.1 M tetrahedra). **(C)** Results of the convergence analysis for the head model with ∼0.9 M tetrahedra, the left M1+DLPFC ROI and the coil positions shown in Fig. 1C.

For the final TMS-optimized head model with ∼1.9 million tetrahedra, we then used a convergence test to assess the numerical accuracy of the simulations. This was done by splitting every edge in the mesh at its midpoint, so that each surface triangle is split into 4 triangles and each tetrahedron is split into 4 tetrahedra and an octahedron. The octahedron was then further split into 4 tetrahedra to create a total of 8 tetrahedra per tetrahedron. This approach preserved the inner angles of the surface triangles and did not degrade surface triangle quality. To increase the quality of the resulting tetrahedra and to reduce their number by a factor of roughly 2, the MMG3D remeshing tools (version 5.7.0) were used. This was repeated three times, creating anatomically identical head models with roughly 1.9 (original), 9.1, 38.3 and 159.1 million tetrahedra. The mean edge lengths are 2.0, 1.2, 0.7 and 0.4 mm for the 4 split levels, covering a similar range as used in the analyses in [3].

As last step, we assessed the differences between simulations with the standard “ernie” and TMS- optimized head meshes. Please note that these differences can stem from differences in the represented anatomical detail and differences in numerical accuracy (due to a varying mesh density). Assessments of the numerical accuracy and of the differences to results for the standard mesh were also performed for the mesh with ∼0.9 million tetrahedra optimized for the left M1+DLPFC ROI, thereby testing 11 coil positions each with 4 orientations above the ROI (Fig. 1C).

### Tests of Speed and Memory Consumption

Initial speed tests were run for the standard “ernie” and the two TMS-optimized meshes on a seasoned laptop (Windows 10, Intel i7-6700HQ with 4 cores, 32GB), using a model of the Magstim 70mm figure-8 coil. Further tests of speed and memory consumption were performed on two standard desktop computers, without (Windows 10, Intel i7-11700, 16 cores, 32GB RAM) and with GPU acceleration (Windows 11, Intel i9-13900KF, 24 cores, 32GB RAM, NVIDIA RTX3060Ti) and for two coil models (Magstim 70mm figure-8, MagVenture Cool D-B80).

For assessing the speed of combined electric field calculations and visualizations in a neuronavigation software on the same computer, a simple TCP/IP protocol between the FEM module and the TMS Navigator software (LOCALITE GmbH, Bonn, Germany) was established utilizing the JSON format. The head model, the tissue conductivities, an initial coil model and a visualization ROI were provided to the FEM module at startup. The communication protocol supports updating the electric field for new coil positions, loading new coil models and visualization ROI(s) during runtime, and dynamically selecting between the loaded coil models and ROIs, respectively.

## Results

### Accuracy of Simulations

Initial tests confirmed that the results were identical up to single precision floating-point accuracy to those obtained with SimNIBS 4.1 and its default iterative preconditioned conjugate gradient solver when using the standard “ernie” head. This confirmed the correctness of the optimized implementation and showed that the choice of solver (direct vs iterative sparse matrix solver) did not affect the simulation outcome.

The difference resulting from cutting the neck at different axial levels is shown in Figure 2A. For the TMS-optimized head mesh used in the main paper, a cutting level of 3 cm below the lowest part of the cranial cavity was chosen (as indicated by the red arrows). This ensured a difference of 2.0±0.9% (mean ± SD) for the electric field magnitude in the bilateral central gray matter ROI. The difference for the electric field vectors was similar.

The convergence test by repeatedly refining the TMS-optimized mesh showed numerical errors of 3.1±0.4% and 4.6±0.5% for the electric field magnitude and vectors for the non-split mesh (Fig. 2B). The difference to the results obtained with the standard “ernie” head was 4.0±0.1% and 6.1±0.1% for the electric field magnitude and vectors. For the electric field calculations in the left M1+DLPFC ROI and the correspondingly optimized head mesh (∼0.9 million tetrahedra), the convergence test showed numerical errors of 4.9±0.3% for the electric field magnitude and 6.7±0.4% for the electric field vectors for the non-split-mesh. The difference to the results obtained with the standard “ernie” head was 3.3±0.1% and 4.9±0.1% for the electric field magnitude and vectors. The slightly lower differences compared to the bilateral gray matter ROI likely stem from the position of the M1+DLPFC ROI in the superior part of the brain, which minimizes the residual effects of the (cut) neck on the current flow.

### Speed and Memory Consumption

Initial speed tests were performed on the same laptop (Windows 10, Intel i7-6700HQ with 4 cores, 32GB) for the standard “ernie” head mesh with ∼4.4 million tetrahedra and the bilateral central gray matter ROI and coil positions shown in Fig. 1B. Update rates for the electric field magnitude were 695.2 ± 45.5 ms (Table 1), and the startup of the FEM module with all pre-calculation steps required ∼173 seconds. Using TMS-optimized meshes improved the update rates further to >3 Hz (bilateral central gray matter ROI) and >10 Hz (left M1+DLPFC ROI).

**Table 1.** Update rates in [ms] (Windows 10, Intel i7-6700HQ with 4 cores, 32GB). Listed are average values (± SD) of 1000 cycles and the Magstim 70mm figure-8 coil.

| Update rates in [ms] | standard mesh (~4.4 M tets) & bilateral gray matter | TMS-optimized mesh (~1.9 M tets) & bilateral gray matter | TMS-optimized mesh (~0.9 M tets) & left M1+DLPFC |
| --- | --- | --- | --- |
| E-field magnitude | 695.2 ± 45.5 | 277.7 ± 20.8 | 87.6 ± 9.5 |
| E-field vectors | 753.6 ± 53.3 | 308.7 ± 25.5 | 89.5 ± 8.9 |

In contrast, SimNIBS 4.1 [3] required 11.3 ± 0.2 s per position to calculate the electric field magnitude at the tetrahedra barycenters for the same head mesh and the PARDISO solver, which is already faster than its default iterative solver. Further ∼5 s per position were spent on creating and saving the visualizations, and determining the electric field magnitude on the bilateral central gray matter ROI via SPR interpolation required additional 49.9 ± 2.0 s per position. These results show that the new implementation achieves a substantial increase in speed even without employing head meshes optimized for TMS calculations on the same hardware.

Update rates for two further computers are listed in Table 2 and range between ∼4.5Hz (bilateral gray matter, no GPU acceleration, electric field calculations and visualizations; Fig. 1D) to ∼27Hz (left M1+DLPFC ROI, GPU acceleration, only electric field calculations). In these cases, the longest startup of the FEM module took ∼45s for the head mesh with ∼1.9 M tetrahedra and the bilateral gray matter ROI. Loading and preparing the left M1+DLPFC ROI during runtime took ∼5s, loading of a coil model <1s. Switching between loaded coil models and/or visualization ROIs incurs no additional update time and is hence feasible for each new coil position. The peak memory consumption of the FEM module stayed below 4GB, which occurred during the startup phase for the head mesh with ∼1.9 M tetrahedra. Afterwards, the average memory consumption was ∼2GB. Both values are uncritical on modern computers.

**Table 2.** Update rates in [ms] (CPU: Windows 10, Intel i7-11700, 16 cores, 32GB RAM; CPU&GPU: Windows 11, Intel i9-13900KF, 24 cores, 32GB RAM, NVIDIA RTX3060Ti). Unless indicated otherwise, the results are for the model of the Magstim 70mm figure-8 coil. Listed are average values (± SD) of 25000 cycles (without visualization) and 1000 cycles (with visualization). As the visualization is currently restricted to the E-field magnitude, update rates were only determined for that case.

| Update rates<br>in [ms] |  | CPU | CPU<br>MagVenture<br>Cool D-B80 | CPU<br>incl.<br>Visualization | CPU&GPU | CPU&GPU<br>Incl.<br>Visualization |
| --- | --- | --- | --- | --- | --- | --- |
| Bilateral<br>gray<br>matter | E-field<br>magnitude | 160.9 ± 11.1 | 163.6 ± 7.7 | 229.5 ± 10.2 | 93.2 ± 9.1 | 136.1 ± 8.4 |
|  | E-field<br>vectors | 169.0 ± 10.9 | 171.8 ± 7.6 | - | 92.8 ± 9.1 | - |
| Left M1-DLPFC | E-field<br>magnitude | 59.4 ± 8.0 | 60.0 ± 7.9 | 83.6 ± 5.4 | 37.5 ± 8.2 | 51.3 ±4.5 |
|  | E-field<br>vectors | 60.1 ± 7.8 | 60.6 ± 7.8 | - | 37.2 ± 8.3 | - |

## Discussion

We introduced and validated calculations of the electric field induced by TMS in the brain that were optimized for rapid repeated evaluations for changing the coil positions or coils. Extending our previous method [3], the approach uses a first-order tetrahedral FEM and super-convergent patch recovery to maintain a good balance between computational accuracy and efficiency.

Repeated evaluations benefit strongly from its update rates of 1 Hz and better, while the required startup time stays moderate with ∼3 min (values for a “standard” 4.4 M tetrahedra mesh). When tested with the same volume conductor model, it achieves identical results as our previous method that has been stringently validated against an analytical solution for a spherical geometry [3] and against a boundary-element method [5].

We also demonstrated that volume conductor models can be optimized for TMS calculations to maintain a similar numerical error (∼3% for the electric field magnitude) with less tetrahedra, compared to “multi-purpose” volume conductor models that are aimed to support both TMS and TES simulations. In combination, the optimized approach and meshes enabled update rates up to 10 Hz for electric field calculations in bilateral gray matter and >20Hz for a large ROI spanning the left M1 and DLPFC. Moderate memory requirements of <4GB for TMS-optimized meshes, good performance even without dedicated GPU hardware and the ability to switch between different coil models and visualization ROIs during runtime support the practical usability of the approach.

### Related work

Several studies presented alternative or complementary approaches to speed up TMS electric field simulations, aimed at various application scenarios. For example, BEM calculations optimized for repeated TMS simulations reached a better numerical accuracy than our approach but had lower update rates and required longer startup times [5]. It thus depends on the specific application whether the numerical accuracy of BEM offers a relevant advantage.

Various surrogate models have been developed to support fast online or real-time electric field simulations. In principle, deep-learning-based (DL) methods can predict the TMS electric field directly from the MRI intensity distribution [2,14], thus resolving the need for creating dedicated volume conductor models (requiring ∼2 hrs for SimNIBS charm). However, they need retraining when changing the coil type or when updating the segmentation and tissue conductivities used during training. In addition, their robustness to contrast changes in the MR images still awaits evaluation. A further DL approach used a three-dimensional tissue conductivity map of the head as input to overcome some of these limitations [11], but in turn needs a prior segmentation of the MR image. The reported calculation speeds ranged from 0.024 s [2] to 1.47 s [11] on dedicated GPU hardware, while accuracy remained lower than that of FEM- or BEM-based methods.

Other approaches gain speed during runtime by pre-calculating “basis sets” from which the TMS electric fields can be approximated [9,10,12,13]. They offer different strengths and weaknesses. Some are limited to flat coil geometries or are restricted to specific coil-scalp distances and tilts (e.g. assuming perfectly tangential coil placements), or coil positions covering only parts of the head. In most cases, the pre-calculations need to be repeated when changing the head model, the coil type or the brain ROI in which the field is evaluated. While some approaches support calculations on the complete cortical surface, the pre-calculations can take several hours [9,13,15] in addition to the time for creating the head mesh. A recent approach resolved most of these limitations [15], but still requires time-consuming pre-calculations for each new patient dataset and GPU acceleration to perform well. Our approach avoids most of these limitations. Depending on the required update rates, it can act as an alternative or can complement the above approaches by helping to strongly shorten the required pre-calculation phases.

### Limitations

The time cost for the automatic head model creation remains as practically relevant drawback of our approach, and future research could aim at accelerating this process. In addition, the JSON-based TCP/IP protocol was used as a straightforward and stable communication solution between the FEM module and the neuronavigation software, but its overhead and time delay could be avoided using more sophisticated approaches based on shared memory. Removing most of the upper neck from the TMS-optimized head models contributed to the overall differences of the electric field magnitudes of ∼4% compared to the results of the standard-resolution head model. When the interest is specifically on inferior coil positions, choosing a lower cut-off level at the cost of increasing the number of tetrahedra would help to improve overall simulation accuracy.

## Acknowledgments

The work was supported by the Innovation Fund Denmark (grant 9068-00025B Precision-BCT). AT was supported by the Lundbeck Foundation (grants R313-2019-622 and R244-2017-196). HRS was supported by a collaborative alliance grant by Lundbeck Foundation (R336-2020-1035). HRS holds a 5-year professorship in precision medicine at the Faculty of Health Sciences and Medicine, University of Copenhagen which is sponsored by the Lundbeck Foundation (Grant Nr. R186-2015-2138). OP is supported by the Lundbeck Foundation (grant R360-2021-395).

